# Pathogenic entero- and salivatypes harbour changes in microbiome virulence and antimicrobial resistance genes with increasing chronic liver disease severity

**DOI:** 10.1101/2023.08.06.552152

**Authors:** Sunjae Lee, Bethlehem Arefaine, Neelu Begum, Marilena Stamouli, Elizabeth Witherden, Merianne Mohamad, Azadeh Harzandi, Ane Zamalloa, Haizhuang Cai, Lindsey A Edwards, Roger Williams, Shilpa Chokshi, Adil Mardinoglu, Gordon Proctor, Debbie L Shawcross, David Moyes, Mathias Uhlen, Saeed Shoaie, Vishal C Patel

## Abstract

**Background & Aims:** Life-threatening complications of cirrhosis are triggered by bacterial infections, with the ever-increasing threat of antimicrobial resistance (AMR). Alterations in the gut microbiome in decompensated cirrhosis (DC) and acute-on-chronic liver failure (ACLF) are recognised to influence clinical outcomes, whilst the role of the oral microbiome is still being explored. Our aims were to simultaneously interrogate the gut and oral micro- and mycobiome in cirrhotic patients, and assess microbial community structure overlap in relation to clinical outcomes, as well as alterations in virulence factors and AMR genes.

**Methods:** 18 healthy controls (HC), 20 stable cirrhotics (SC), 50 DC, 18 ACLF and 15 with non-liver sepsis (NLS) *i.e.* severe infection but without cirrhosis, were recruited at a tertiary liver centre. Shotgun metagenomic sequencing was undertaken from saliva (S) and faecal (F) samples (paired where possible). ‘Salivatypes’ and ‘enterotypes’ based on clustering of genera were calculated and compared in relation to cirrhosis severity and in relation to specific clinical parameters. Virulence and antimicrobial resistance genes (ARGs) were evaluated in both oral and gut niches, and distinct resistotypes identified.

**Results:** Specific saliva- and enterotypes revealed a greater proportion of pathobionts with concomitant reduction in autochthonous genera with increasing cirrhosis severity, and in those with hyperammonemia. Overlap between oral and gut microbiome communities was observed and was significantly higher in DC and ACLF *vs* SC and HCs, independent of antimicrobial, beta-blocker and acid suppressant use. Two distinct gut microbiome clusters [ENT2/ENT3] harboured genes encoding for the phosphoenolpyruvate:sugar phosphotransferase system (PTS) system and other virulence factors in patients with DC and ACLF. Substantial numbers of ARGs (oral: 1,218 and gut: 672) were detected with 575 ARGs common to both sites. The cirrhosis resistome was significantly different to HCs, with three and four resistotypes identified for the oral and gut microbiome, respectively.

**Discussion:** Oral and gut microbiome profiles differ significantly with increasing severity of cirrhosis, with progressive dominance of pathobionts and loss of commensals. DC and ACLF have significantly worse microbial diversity than NLS, despite similar antimicrobial exposure, supporting the additive patho-biological effect of cirrhosis. The degree of microbial community overlap between sites, frequency of virulence factors and presence of ARGs, all increment significantly with hepatic decompensation. These alterations may predispose to higher infection risk, poorer response to antimicrobial therapy and worsening outcomes, and provide the rationale for developing non-antibiotic-dependent microbiome-modulating therapies.

## Introduction

One in five hospitalised patients with cirrhosis die (*1*). The prospective multicentre PREDICT study showed that almost all patients with acute decompensation with and without the development of acute-on-chronic liver failure (ACLF) had proven bacterial infections (BIs) as a precipitant (*2*). BIs caused by extensively drug-resistant (XDR) organisms are associated with the highest risk of developing (multi-)organ failure and account for nearly half of cases globally (*3*). BIs in patients with decompensated cirrhosis (DC) typically result from breaches in innate immune barriers and inadequate clearance by immune cells (*4*).

Antimicrobial therapy therefore forms the cornerstone of treatment in cirrhosis, both for acute BIs and also as prophylaxis against infection-driven complications (*5*). This is however mired with challenges due to diagnostic delays and uncertainties, and an increasing frequency of multidrug resistance organisms (MDRO) (*6, 7*). High levels of antimicrobial resistance (AMR) deleteriously affecting the outcome of treatment with antibacterial agents in cirrhosis is increasingly a cause for concern in Europe (*8*) and worldwide (*9*). This is particularly worrying in cirrhotic patients who have a heightened susceptibility to BIs as a consequence of a distinctive spectrum of immune alterations, termed cirrhosis-associated immune dysfunction (CAID) (*10*).

Alterations in the gut microbiome in DC and ACLF – the latter being characterised by organ failure, critical illness and very high short-term mortality – are recognised as being pivotal in influencing clinical outcomes (*11, 12*) and mechanistically contributing to hepatic decompensation (*13*). This so-called gut ‘dysbiosis’ is causally linked to MDRO infections, due to intestinal barrier and mucosal immune homoeostatic failures, enhanced pathobiological translocation of microbes and their metabolites (including toxins) and deficits in host-microbiome compartmentalisation (*14*). Multiple studies over the past decade have reported alterations in the gut microbiome in cirrhosis, and its role in hepatic decompensation (*13*). Alterations of individual genera in the gut have been associated with cirrhosis progression, namely *Enterococcacae* and *Enterobacterceae* (*15*), as well as changes in the oral microbiome (*16*). The expansion in knowledge has largely been driven by next generation sequencing (NGS) of faecal samples, as a surrogate for the intestinal niche, largely due to the non-invasiveness of material acquisition. The majority of studies to date on human microbiome abnormalities in cirrhosis have employed less phylogenetically resolving 16S rRNA gene sequencing approaches instead of more advanced shotgun metagenomics.

The functional relevance of gut and oral microbial alterations is being recognised as more clinically and mechanistically purposeful than unilaterally describing varying phylogenetic levels of compositional changes (*17*), and is increasingly linked to metaproteomic, metatranscriptomic and metabolomic profiling (*18–20*). The gut microbiome in cirrhotic patients compared to healthy controls in a seminal study reported higher levels of *Streptococcus* and *Veillonella* species - species usually detected in the oral cavity - present in cirrhotic faeces (*21*). Comparison of faecal bacterial species in these cirrhotic patients with those known to be commensal within the oral cavity and gut of healthy individuals demonstrated a partial similarity to both oral and ileal bacteria of the healthy individuals. Over 75,000 microbial genes differed in this study between cirrhotic and healthy individuals, and over 50% taxonomically assigned bacterial species were of oral origin suggested by the authors as an ‘invasion’ of the distal gut from the mouth in cirrhosis.

This hypothesis that oral microbes can extend into and/or invade the lower intestine may be a consequence of changes in intestinal pH and/or bile acid dysregulation (*22, 23*) that occur in advanced cirrhosis. The relocation of oral microbes into the distal intestine may also be related to an epiphenomenon predisposed by impaired gastric acid and bile secretion that is prevalent in cirrhosis. This increase in gastric and intestinal pH is further exacerbated by the use of proton pump inhibitor (PPI) therapies widely prescribed in cirrhotics, and PPI use has been reported as altering gut microbiome community structures in large healthy cohorts (*24, 25*). A significantly elevated relative abundance of *Streptococcaceae*, which are typically limited to the oral cavity, was detected in faeces prior to and following PPI therapy in patients with cirrhosis (*26*) implying that modifications of gastric and bile acid chemical barriers facilitate this migration and outgrowth of oral commensals within the distal intestine. In addition, conventional gut-targeting therapies established for the prevention of hepatic encephalopathy (HE) in DC such as rifaximin-α have been shown to impact on not only the gut microbiome but also suppressing oral bacteria (*27*). These oral species have putative functions related to intestinal mucus degradation, so that a reduction in these reaching the gut promotes gut barrier integrity, and in doing so, ameliorates HE symptoms.

A recent study reported marked compositional alterations in the faecal microbiome that parallel disease stages with maximal changes in ACLF, in varying stages of cirrhosis, even after adjustment for antibiotic therapy (*28*). Functional insights enabled by analysis of shotgun metagenomic sequences demonstrated that cirrhotic patients had enriched pathogenic pathways related to ethanol production, gamma-aminobutyric acid metabolism, and endotoxin biosynthesis. These changes were positively associated with complications of cirrhosis and 3-month survival. However, the oral microbiome was not profiled so comparisons to the faecal microbiome, as well as an interrogation of oral microbial functional pathways, was not possible. Analysis of the fungal microbiome - termed ‘mycobiome’ and antimicrobial resistance genes (ARGs) were also lacking, all of which are of relevance in the pathobiological ‘oral-gut-liver axis’ in cirrhosis.

The gut is a reservoir for ARGs with disruption of the microbiome leading to colonisation by pathogenic organisms (*29*). Bacterial species can acquire resistance genes through horizontal gene transfer and the high-density communities found in the gut give rise to a wide distribution of ARGs (*30*). The burden of ARGs within the gut microbiome reservoir is a functional threat in cases of dysbiosis (*31*). A better understanding of the ARGs harboured by the oral and gut microbiome in cirrhosis is critical, given the escalating rates of AMR, the contribution of the microbiome to heightened infection risk, and data demonstrating the strong association of MDRO BIs on mortality, especially in ACLF (*3*).

This study aimed to simultaneously interrogate the gut and oral microbiome and mycobiome utilising deep shotgun metagenomic sequencing of faecal and saliva samples, respectively, in well-phenotyped cirrhotic patients of varying disease severities, in comparison with healthy and positive disease controls. Our objectives were to assess (i) the degree of overlap and alterations between oral and gut microbiome community structures, (ii) virulence factors and ARG carriage, and (iii) crucially how these evolve with increasing severity of cirrhosis and the impact of organ failure and critical illness. Finally, we provide novelty in exploring how these changes relate to clinically relevant parameters and endpoints at different stages of cirrhosis.

## Materials & Methods

### Study participants & biological sampling

Patients were consecutively recruited at King’s College Hospital after admission to the ward or from the hepatology out-patient clinic. The study was granted ethics approval by the national research ethics committee (12/LO/1417) and the local research and development department (KCH12-126) and performed conforming to the Declaration of Helsinki. Patient participants, or their family nominee as consultees in the case of lack of capacity, provided written informed consent within 48 hours of presentation. Patients were managed according to standard evidence-based protocols and guidelines (*32*).

Patient participants were stratified into and phenotyped according to clinically relevant groups based on the severity and time course of their underlying cirrhosis, degree of stability and hepatic decompensation, and presence and extent of hepatic and extra-hepatic organ failure at the time of sampling. These groups were stable cirrhosis (n=20), acutely decompensated cirrhosis (AD) (n=50) and acute-on-chronic liver failure (ACLF) (n=18), with a separately recruited healthy participant control cohort (n=40). AD was defined by the acute development of 1 or more major complications of cirrhosis, including ascites, hepatic encephalopathy, variceal haemorrhage, and bacterial infection. ACLF was defined and graded according to the number of organ failures in concordance with criteria reported in the CANONIC study (*33, 34*). Main exclusion criteria included pregnancy, hepatic or non-hepatic malignancy, pre-existing immunosuppressive states, replicating HBV/HCV/HIV infection, and known IBD.

Demographic, clinical, and biochemical metadata were collected at the time of biological sampling. Standard clinical composite scores used for risk stratification and prognostication included the Child-Pugh score (*35*), model for end-stage liver disease (MELD) (*36*), United Kingdom model for end-stage liver disease (UKELD) (*37*), Chronic Liver Failure Consortium-acute decompensation (CLIF-C AD) (*38*) and Sequential Organ Failure Assessment (SOFA) (*39*).

For patients with sepsis without CLD (non-liver sepsis (NLS), n=15), the diagnosis of sepsis was based on the Sepsis-3 criteria (*39*) in which life-threatening organ dysfunction caused by a dysregulated host response to infection was evident, with organ dysfunction defined by an increase in the sequential (sepsis-related) organ failure assessment (SOFA) score of 2 points or more. The absence of CLD in this patient group was determined by a combined assessment of clinical history with biochemical and radiological parameters.

Healthy controls aged >18 years were recruited to establish reference values for the various assays performed. Exclusion criteria for healthy controls were body mass index <18 or >27; pregnancy or active breastfeeding, a personal history of thrombotic or liver disease; chronic medical conditions requiring regular primary or secondary care review such as inflammatory bowel disease, and/or prescribed pharmacotherapies

### Faecal sample acquisition

Faecal samples were obtained within 48 hours of admission to hospital and collected into non-treated sterile universal tubes (Alpha LaboratoriesTM), without any additives. Faecal samples were kept at 4°C without any preservative and within 2 hours were homogenised, pre-weighed into 200mg aliquots in Fastprep tubes (MP BiomedicalsTM), for storage at −80°C for subsequent DNA extraction.

### Saliva sample acquisition

Saliva samples were obtained within 48 hours of admission to hospital and collected into non-treated sterile universal tubes (Alpha Laboratories^TM^), without any additives. A controlled passive ‘drool’ was performed by the study participant into a universal container repeatedly until at least 6mL of saliva was obtained. For patients that were intubated for mechanical ventilation, oro-pharyngeal suctioning of accumulating oral secretions were obtained. Saliva samples were kept at 4°C without any preservative and within 2 hours were homogenised, and measured into 1mL aliquots using sterile wide bore pipettes in Fastprep tubes (MP Biomedicals^TM^), which were then centrifuged at 17,000 g for 10 minutes. The saliva supernatant was removed and stored separately whilst the remaining pellet was stored at −80°C for subsequent DNA extraction.

### DNA extraction from faecal and saliva samples

A two-day protocol adapted from the International Human Microbiome Standards (IHMS) (*40, 41*) was used to extract DNA from both stored faecal and saliva pelleted samples. For faeces, a 200mg pre-weighed and homogenised aliquot was used and for saliva, a post-centrifugation pellet was used. Please refer to the Supplementary section for further details on extraction protocol.

### Library preparation

Illumina TruSeq DNA PCR-free library preparation (Illumina Cat no: 20015963, Illumina, USA) was used to generate high quality DNA sequencing libraries, adapted for automation for the Aglient NGS Bravo workstation (Agilent Technologies, USA) in a 96-well plate format. TruSeq PCR-free libraries have better coverage of GC-rich regions compared to PCR-based methods and the reads are more evenly distributed over the genome. The following workflow was adopted: quality checks on the DNA; fragmentation of DNA to obtain 350 bp fragments; library preparation, which consists of bead clean-up of the fragments, phosphorylation, bead-based removal of long and short fragments, addition of a single A nucleotide to the 3’ end (A-tailing), ligation of adapter sequences (120 bp long), bead-based purification of samples to remove non-ligated adapter oligonucleotides; qPCR of prepared libraries to assess concentration; pooling at equimolar concentrations; sequencing.

DNA quantity and quality were assessed using the Qubit fluorometer (ThermoFisher Scientific, USA) with the dsDNA HS Assay Kit, and the TapeStation [using the genomic DNA ScreenTape and Reagents], respectively. The optimal amount of DNA for this method was 1ug. DNA samples were diluted (where applicable) and fragmented using the Covaris S220 system (Covaris, USA) using the 350 bp setting. The subsequent steps of library preparation were carried out on the Agilent NGS Bravo workstation in 96-well plates adapted for the TruSeq PCR-free Sample Preparation protocol by Illumina.

DNA was purified by magnetic beads on the Agencourt AMPure XP (Beckman Coulter) and subsequently used to generate standard barcoded libraries by the Biomek FXp (Beckman Coulter) liquid handler using the Fragment Library Preparation 5500 Series SOLiD™ System (ThermoFisher Scientific) and the IonXpress Plus gDNA Fragment Library Preparation Kit (ThermoFisher Scientific). SPRI beads (Beckman Coulter) were used for clean-up steps of 350 bp fragments; the ligation step added 120 bp to each fragment yielding 470 bp fragments (350 +120 bp). Obtained libraries were either stored at 4°C for up to two days or at −20°C longer-term. The IonXpress Barcode Adapters 1-96 Kit (ThermoFisher Scientific) was used to barcode all libraries. The volume of libraries was estimated by pipetting and the 2100 Bioanalyzer (Agilent, USA) was used to perform quality control of the amplified libraries, together with the High Sensitivity DNA kit (Agilent, USA). Concentrations of the libraries were measured by qPCR using the Kapa library quantification kit (Roche). Libraries with a final concentration of >2 nM were deemed to be suitable for sequencing.

### Whole-genome shotgun metagenome sequencing

TruSeq PCR-free libraries are well suited for deep sequencing to achieve high-coverage genomes. The library preparation method shows a significantly better coverage of GC-rich regions compared to PCR-based methods and the reads are more evenly distributed over the genome. Libraries with an average size of 350 base pairs were validated, normalised, pooled and loaded onto NovaSeq S4 flowcells (Illumina, USA) and sequenced on a NovaSeq 6000 (Illumina, USA). Samples were in 3 lanes S4-300, generating a minimum of 40 million reads per sample. Raw sequencing reads were filtered for High Quality (HQ) reads to a minimum of 20 million per sample, before cleaning to remove possible contaminating human and food-associated reads. This was achieved by mapping the HQ reads to the human reference genome (GRCh39), food-related genomes, Bos Taurus (May 2014 version), and Arabidopsis thaliana (May 2014 version). The resulting HQ-cleaned reads were then mapped and counted using the METEOR pipeline (https://forgemia.inra.fr/metagenopolis/meteor).

### Statistical analysis of clinical data

Continuous data were tested for normality using the D’Agostino Pearson test. Comparisons between two or more groups was done by Student’s t test (or Analysis of Variance) and Mann-Whitney U test (or Kruskall Wallis) for normally and non-normally distributed data, respectively. Normally distributed data (*) are presented as mean ± standard deviation (SD) and non-normally distributed data are presented as median (interquartile range). Comparison between categorical data was done by χ2 test or Fisher’s exact test for small sample sizes and data are presented as number (%). Significance was defined at a 95% level and all p values were 2-tailed. Analyses were undertaken utilising IBM SPSS (version 27) and GraphPad Prism (version 9.5.1).

### Microbiome taxonomic profiling

We mapped faecal and saliva metagenomics samples to gene catalogues of gut and oral metagenomes using METEOR program (*42, 43*) (https://forgemia.inra.fr/metagenopolis/meteor). Normalised gene counts of multiple mapped reads by their numbers, we estimated gene counts of gene catalogues of faecal and saliva metagenomic samples. To reduce the sample variability by sequencing depths, gene counts were rarefied into the same 10 million reads per sample and normalised based on their gene lengths and total counts. Normalised gene counts were then used for the quantification of abundances of MSPs by the median value of 25 marker genes representing robust centroids of MSPs (*27*), which was performed by R momr (metaOMineR) package. We estimated the alpha and beta diversity of faecal and saliva samples by Shannon index and Bray-Curtis distance using R vegan and ape packages. Enterotypes and salivatypes were identified by an unsupervised clustering method, Dirichlet multinomial mixture model, which was implemented in R *dmn* package.

For the detection of fungi populations from shotgun metagenomics, fungal gene information from 2,339 fungal genomes from NCBI was downloaded (as of May 2017). Genemark-ES program was used to identify genes from unannotated genomes and filtered out genomes without available gene annotations. After performing CD-HIT-EST to remove redundant genes and total 2,440,644 fungal genes were used to map shotgun metagenomics data. Using METEOR (https://github.com/sysbiomelab/meteor_pipeline) (*44*), we generated gene counts of all fungal genes and normalised them by lengths and multiple mapping. Based on medial values genes per each fungal species, we estimated the abundance of fungal species.

### Functional microbiome analysis

Amino acid sequences of gene catalogues of gut and oral metagenomes were mapped against KEGG orthology (KO) sequences from the KEGG database (version 82) using DIAMOND (ver 0.9.22.123) (*45*). KEGG module information was downloaded from the KEGG database, and we filtered out modules existing only in Eukaryotes. We summed up the normalised gene counts per genes of corresponding KO and compared their abundances between different sample groups by Wilcoxon rank sum tests and performed hypergeometric tests to identify enriched KEGG modules among significantly differential KOs and enrichment tests.

### Antimicrobial resistance gene analysis

Gene catalogues of gut and oral metagenome were mapped against antimicrobial resistance genes, which were downloaded from the CARD database (version 3.0.0), by BLASTP supported from Resistome Gene Identifier (RGI) application (*46*). For the analysis of antimicrobial gene abundances, we summed up the normalised gene counts per each antimicrobial gene *(i.e*., anti-microbial gene ontology terms defined in the CARD database) and analysed the difference of abundance between different sample groups (*e.g.* different enterotype and salivatype) by Wilcoxon rank sum tests. Drug class information of corresponding antimicrobial genes were also obtained from the CARD database.

## Results

### Participant characteristics

This study included 66 patients with cirrhosis (18-75 years of age) classified according to EASL-CLIF consortium criteria as stable cirrhosis (SC, n=26), decompensated cirrhosis (DC, n=46) and acute-on-chronic liver failure (ACLF, n=14) and a cohort of 15 gender-matched healthy controls (HC). Uniquely for this type of cirrhosis-focused study, a cohort of 14 patients with sepsis but without cirrhosis (non-liver septic, NLS) was included, acting as a ‘positive disease control group’ to enable comparisons of patients with an active infection undergoing similar clinical treatments including antimicrobial exposure, but without the pathobiological effects of underlying chronic liver disease.

Table 1 summarises demographic, clinical and biochemical characteristics of the recruited patients. Cirrhosis and NLS patients were older than HC. Predominant aetiologies of cirrhosis included alcohol-related liver disease (ArLD) (SC/DC/ACLF: 50%/63%/71.4%) and non-alcohol-related metabolic-associated steatotic liver disease (MASLD) (SC/DC/ACLF: 7.7%/17.4%/7.1%), respectively. DC and ACLF patients presented with ascites (76.1%/71.4%) and hepatic encephalopathy (8.7%/42.9%) as the predominant manifestation of hepatic decompensation, respectively. None of the AD and ACLF patients had experienced a variceal haemorrhage in the 7 days prior to recruitment nor had developed prior spontaneous bacterial peritonitis during that admission. DC, ACLF and NLS patients were more frequently receiving antibiotics (71.7%/100%/100%, respectively) compared to SC (26.9%) at the time of sampling, reflecting conventional clinical practice. There was no difference in use of rifaximin-α across the three cirrhosis groups, nor in proton pump inhibitor and H2 antagonist use including the NLS group. DC and ACLF patients were more likely to be treated with lactulose and non-selective beta-blockers than SC and NLS groups. Haematological, biochemical and disease severity and prognostic composite scores followed expected patterns for the varying cirrhosis cohorts. Mortality rates at 30-days and 1-year were significantly higher, as expected, when comparing SC to DC and ACLF groups. Almost a fifth and approaching one third of patients with DC and ACLF, respectively, died, whilst approximately a quarter of SC and ACLF and over a third of DC patients were transplanted, over the 12 month follow-up period.

**Table 1.**
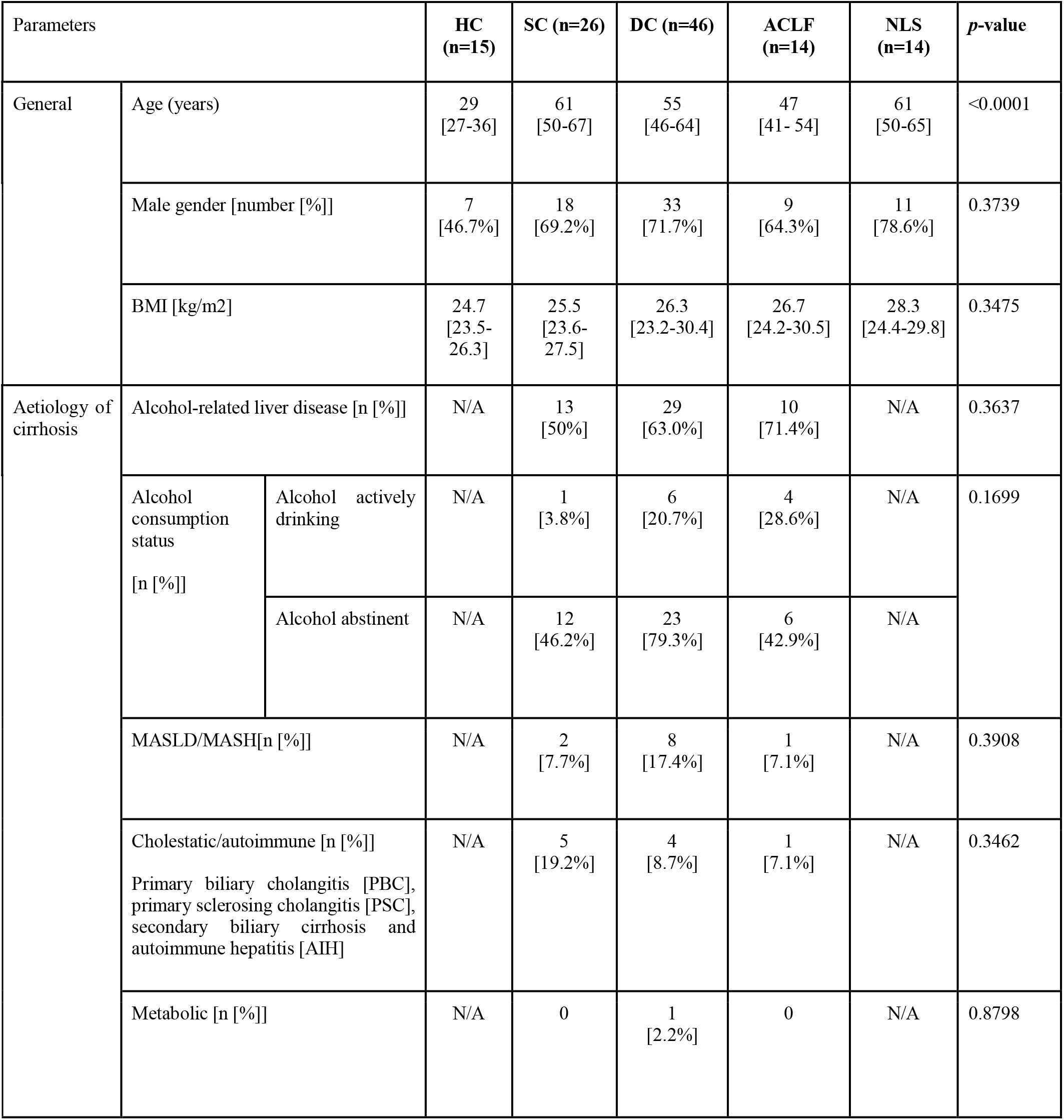
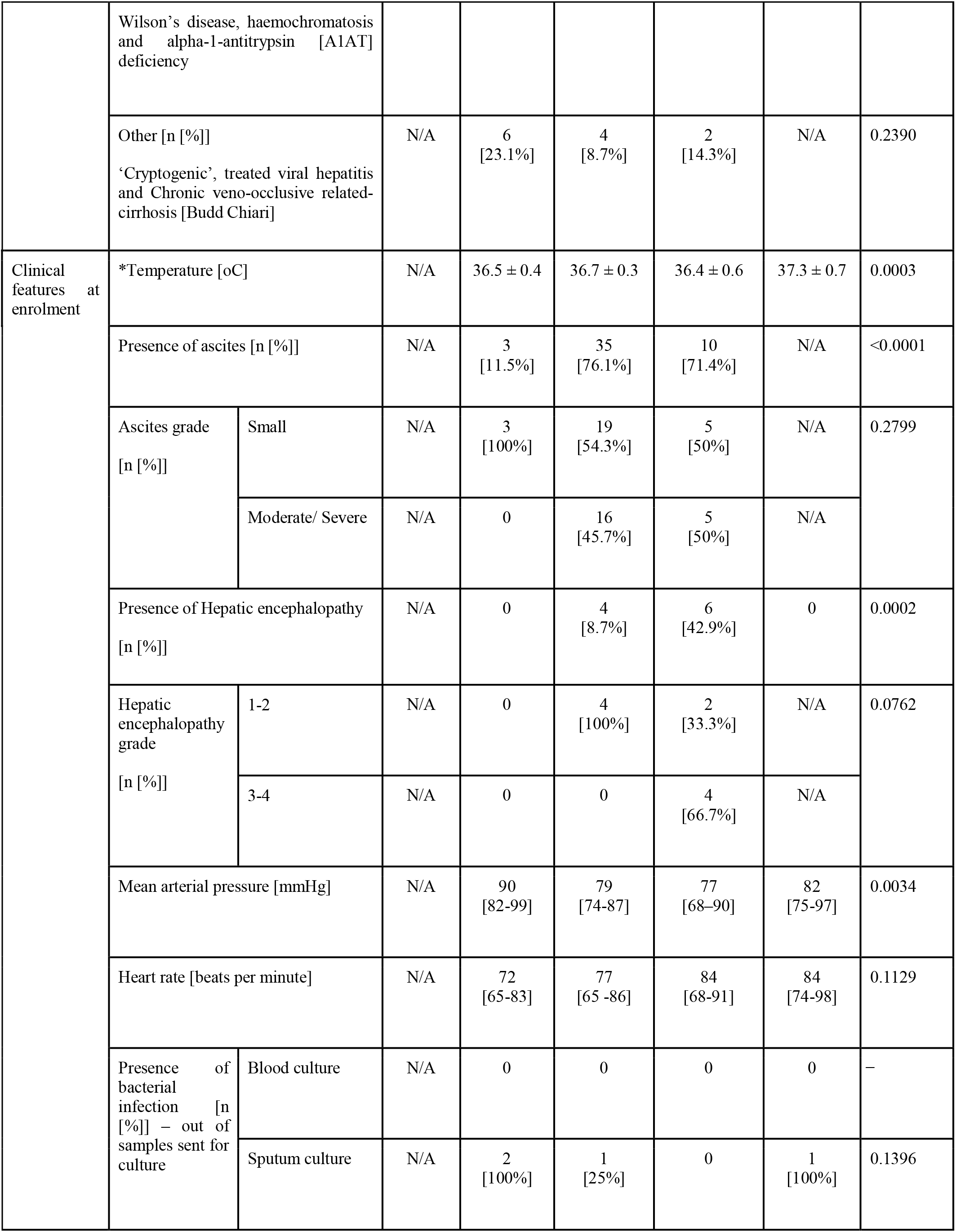
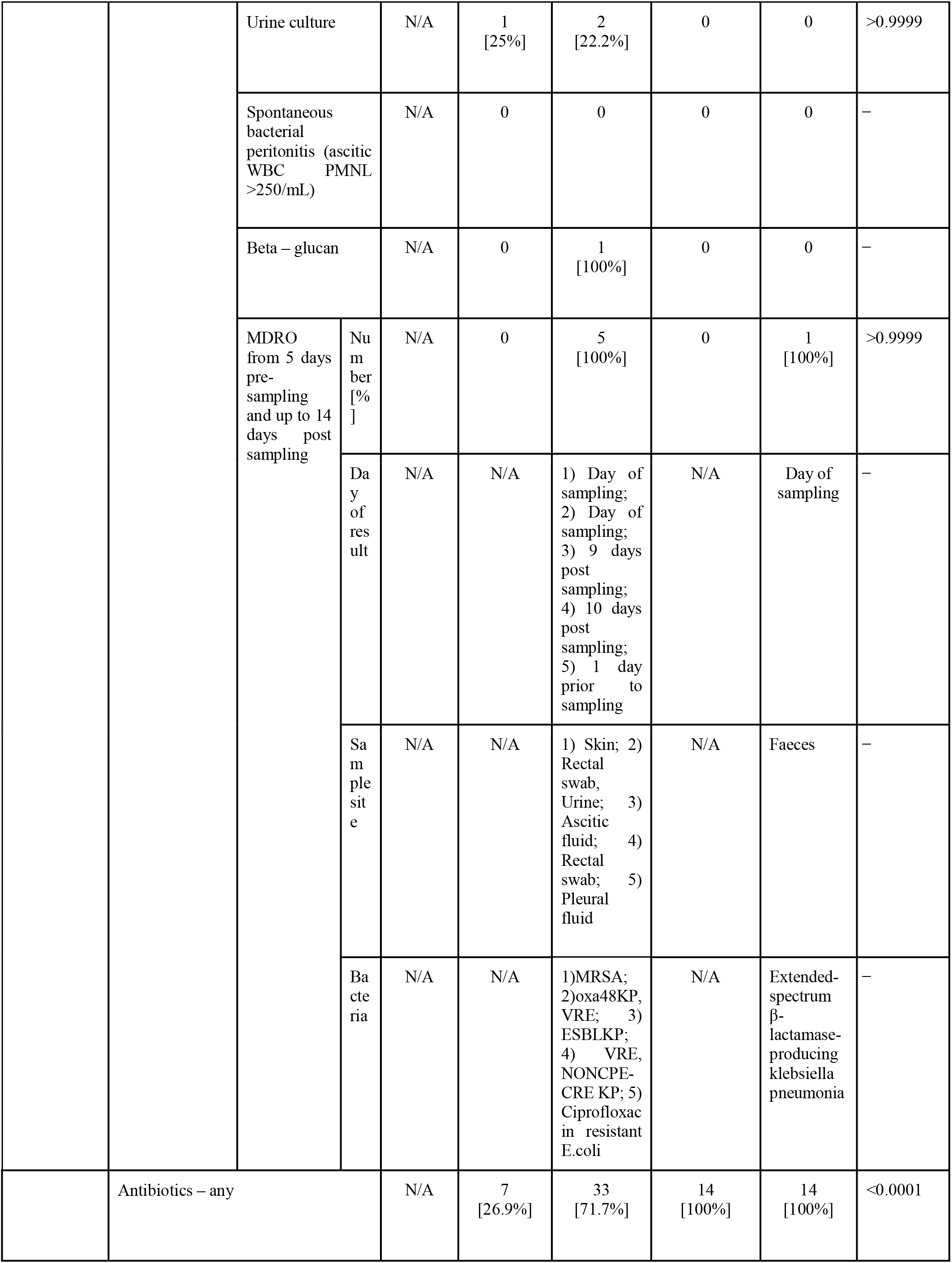
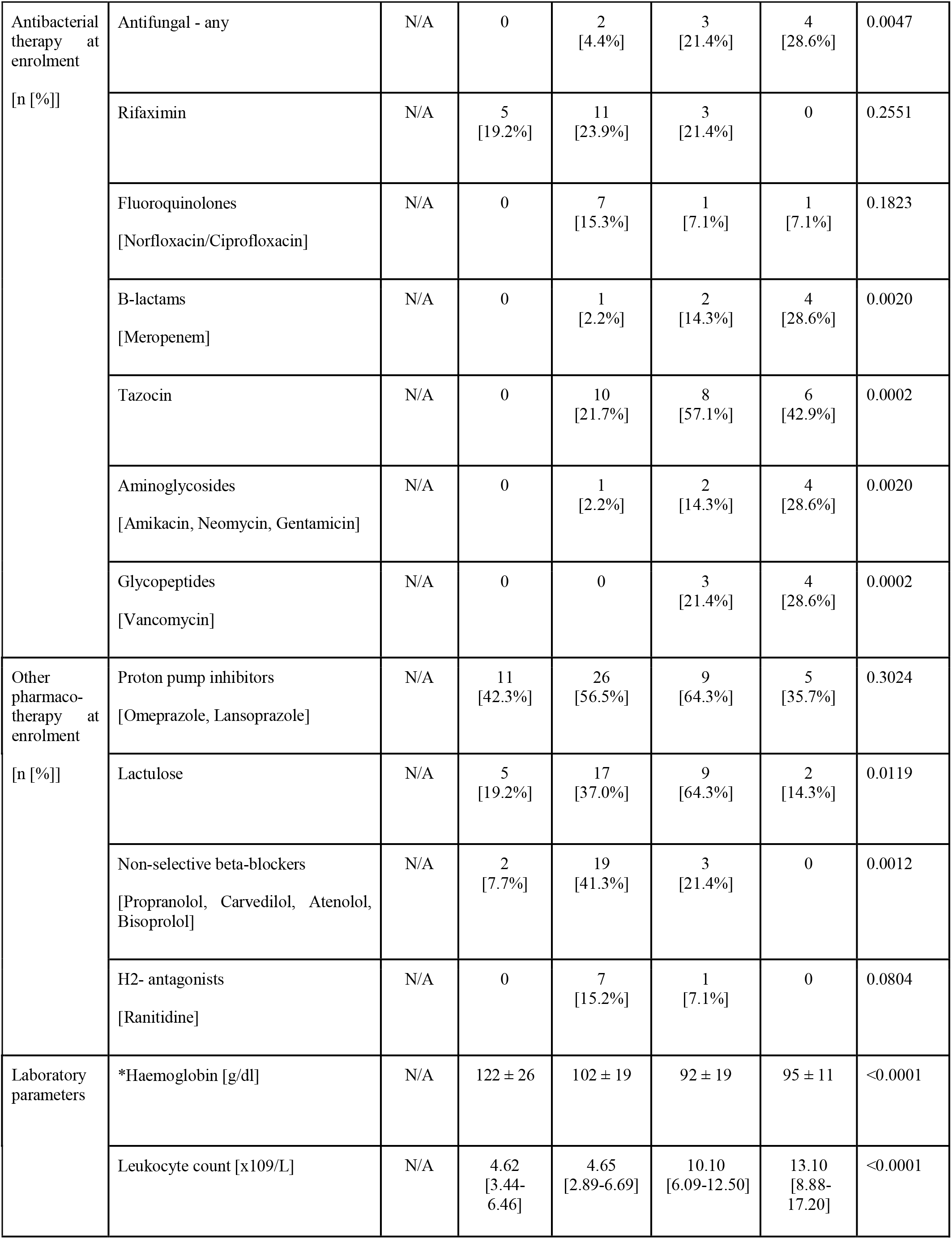
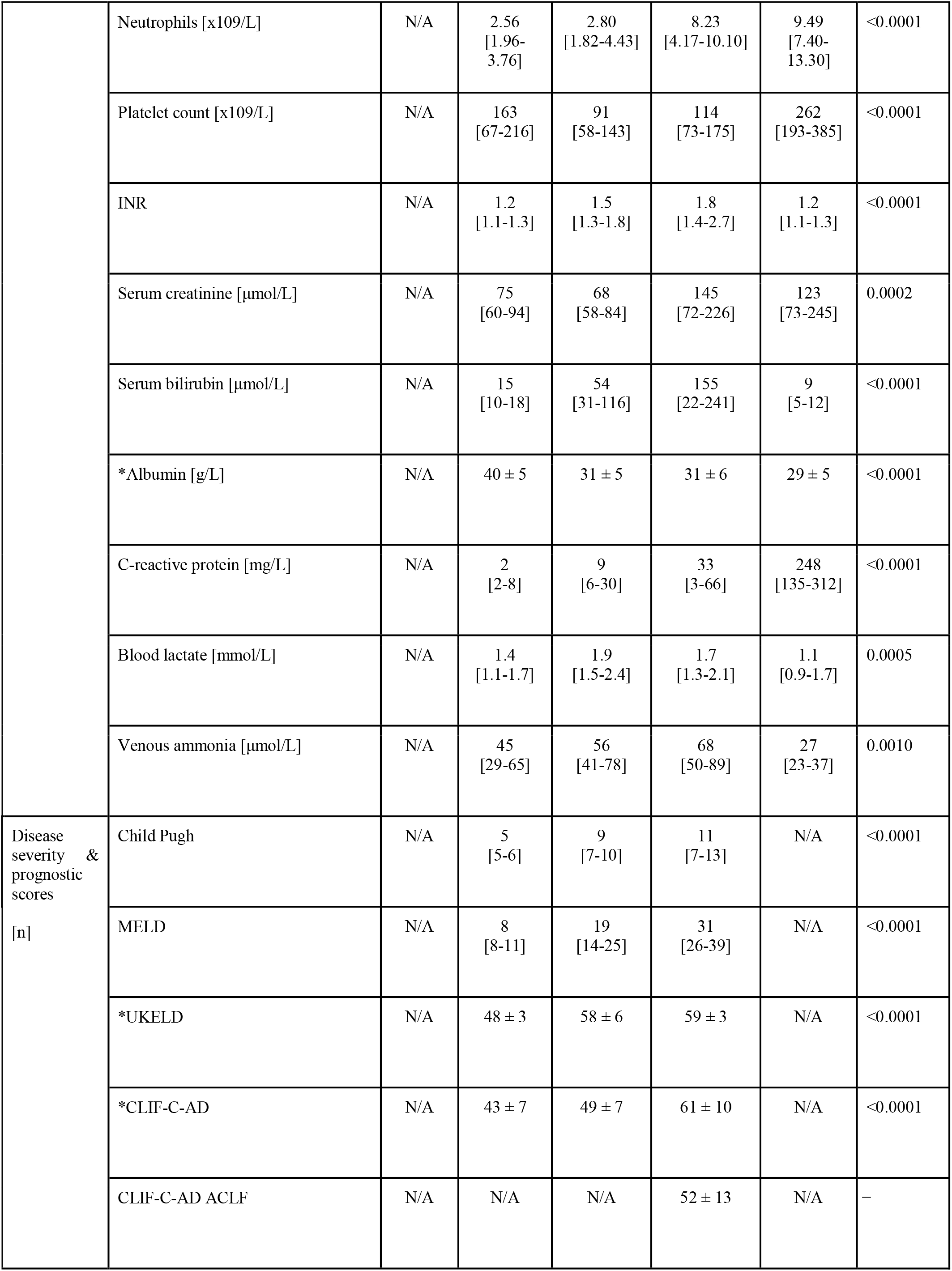
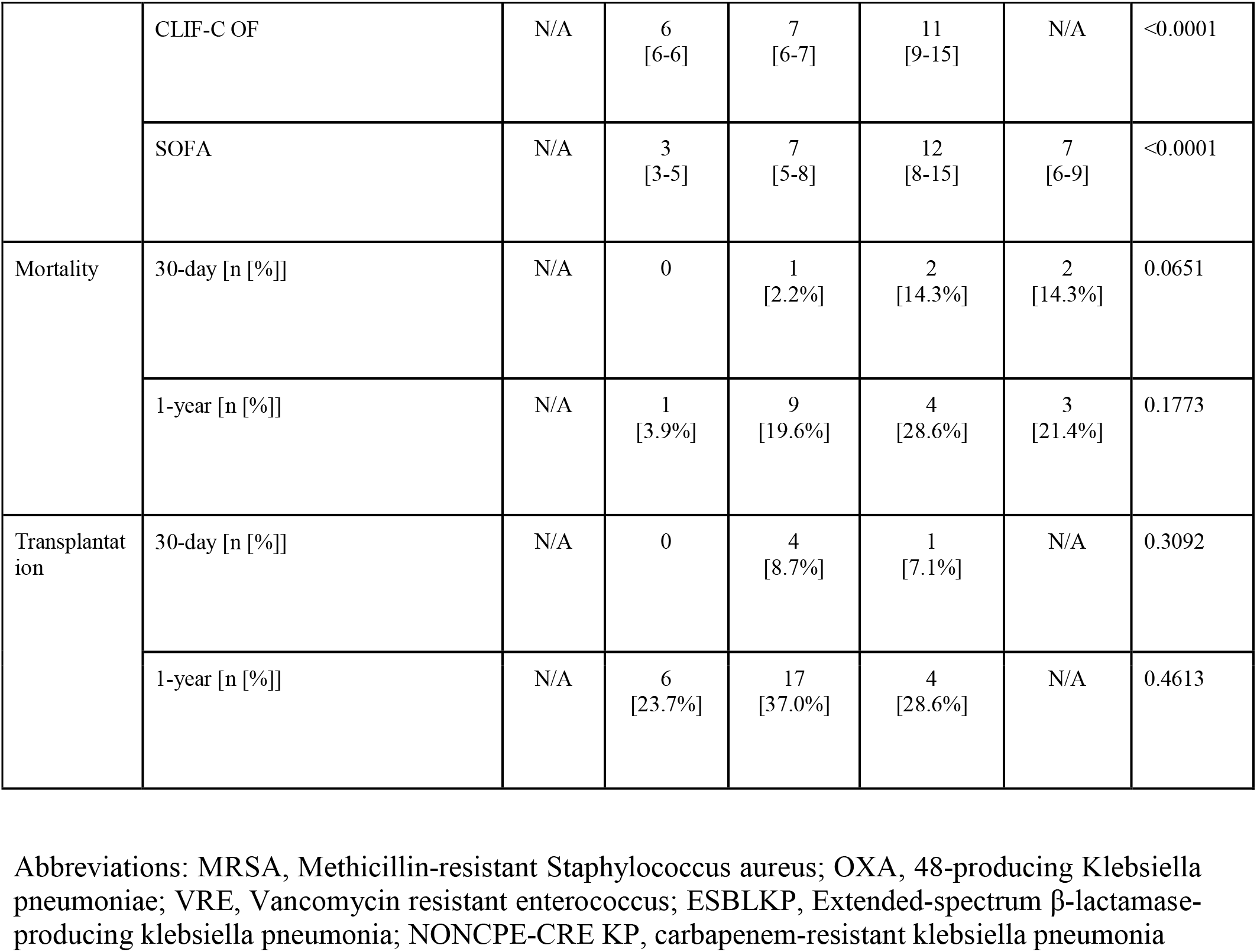
Summary of clinical characteristics of study groups (normally distributed values are denoted with(*) and are presented as mean ± standard deviation (SD); non-normally distributed values are presented as median (interquartile range).). BMI records were missing for 8 HC subjects; platelet count records were missed for 1 DC subject; CRP record was missing for 1 SC subject; blood lactate records were missed for 3 SC subjects, 10 DC subjects and 1 ACLF subjects; venous ammonia records were missed for 6 DC subjects, 2 ACLF subjects, and 5 SEP subjects.

### Compositional alterations in oral and gut microbiome communities in cirrhosis

81 and 66 patients with cirrhosis of varying severities, 11 and 7 with NLS and 15 and 13 HCs underwent biological sampling for faeces and saliva, respectively. We first generated deep-sequenced shotgun metagenomics data of the faecal and saliva samples (minimum 20 million high-quality (HQ) clean reads). We aligned the sequenced reads to the gene catalogues of oral (*42*) and gut microbiome (*43*) and normalised the gene counts after rarefying aligned reads to the same sequencing depth. Using metagenomics species pan-genomes (MSPs) (*47*) as references, we calculated the abundance of microbial species within the faecal and saliva samples (**See Methods**).

We first evaluated the diversity of the gut and oral microbiome by the Shannon index, an alpha-diversity measure of richness and diversity within each sample (**Figures 1A and 1B**). We found both the gut and oral microbiome communities showed significant reductions in alpha-diversity (Wilcoxon rank sum tests p-values <0.05) in keeping with lower microbial community richness and diversity, with increased cirrhosis severity and hepatic decompensation. Next, we checked contrasted taxa at gut and oral sites (*i.e*. at a family level) between the different cirrhosis and control cohorts. Here, particular patterns emerged in relation to the study groups, in particular between the various cirrhosis groups, differentiated by increasing disease severity. For example, we found that opportunistic pathobionts, including *Enterococcaceae* and *Enterobacteriaceae*, and bacterial families strongly associated with cirrhosis, including *Veillonellaceae* and *Streptococcaceae*, significantly increased in both niches as cirrhosis severity increased (**Figure 1C and D**). Conversely, a decreasing relative abundance in taxa conventionally classified as autochthonous or commensal was observed. These gut bacterial families, including *Oscillospiraceae* and *Ruminococcaceae*, and oral commensals, including *Neisseriaceae* and *Prevotellaceae*, decreased as cirrhosis severity increased. Thus an increasing proportion of pathobionts with a relative reduction in commensal bacteria drove a significant alteration in overall microbial community structures affecting both the gut and oral niches simultaneously.

**Figure 1.**
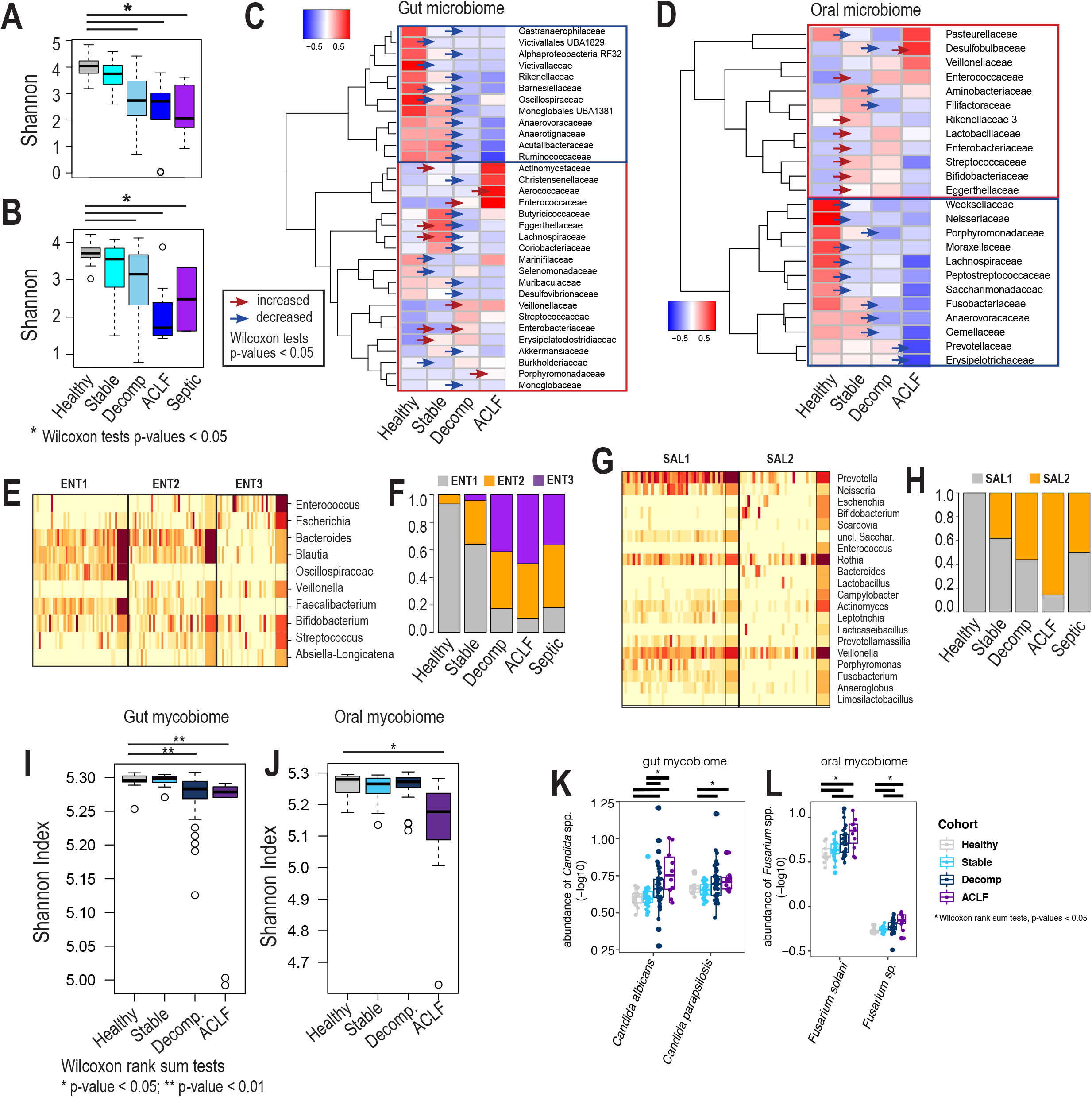
microbial diversity and community structures changed by disease severity. A-B, alpha-diversity of (A) gut and (B) oral microbiome by different disease severity. Significant reduction of alpha-diversity has been observed in more severe groups (Wilcoxon rank sum tests p-values < 0.05). C-D, significantly contrasted family of (C) gut and (D) oral microbiome between adjacent stages of liver cirrhosis. Families with significant changes by stages (Wilcoxon rank sum tests p-values < 0.05) were denoted with red and blue arrows (increase and decrease, respectively). E-H, identification of microbial community structure by unsupervised clustering method – enterotype and saliva-type. We found optimal three microbial clusters from gut microbiome of different disease severities, named ENT1, ENT2 and ENT3 (E), and two microbial clusters from oral microbiome of different disease severities, named SAL1 and SAL2 (F). We found changed fraction of different (F) enterotypes and (H) salivatypes by disease severities. For example, fractions of ENT2/ENT3 and SAL2 were increased by severity, whereas those of ENT1 and SAL1 were decreased. (I) Gut and (J) oral mycobiome diversity was measured with Shannon index and compared with different severity groups (Wilcoxon rank sum tests). Based on Wilcoxon rank sum tests of fungal strains from different severity groups, we identified the alteration of (K) gut fungi and (L) oral fungi (Wilcoxon rank sum tests, p-values < 0.01). For example, candida spp. were increased in the gut mycobiome with worsening cirrhosis severity, whereas *Fusarium* spp was increased in the oral mycobiome.

Next, we identified the hidden community structures of the gut and oral microbiome by an unsupervised clustering method, called a Dirichlet multinomial mixture modelling (see **Methods**). In short, unsupervised clustering identifies distinct clusters within metagenomically analysed samples by maximal separations using the Expectation-Maximisation (EM) process. This approach enabled the identification of three distinct clusters within the gut microbiome (ENT1/2/3; **Figure 1E**) and two distinct clusters within the oral microbiome (SAL1/2; **Figure 1G**), denoted as ‘enterotypes’ and ‘salivatypes’, respectively.

We found enrichment of known genera for enterotypes, such as *Bacteroide*s in ENT1 and ENT2. However, we also found that pathobionts such as *Enterococcus* were dominant in certain enterotypes (ENT2 and ENT3; **Figure 1F**). Notably, bacteria which are usually commensal within the oral niche, such as *Veillonella* and *Streptococcus*, were also enriched in ENT2 and ENT3, in keeping with the transfer of these bacteria from the oral cavity into the lower gut. This replicates the microbiome patterns previously reported in another DC trial setting comparing oral and faecal bacterial communities (*27*). Notably, the relative proportion of these oral commensals being detected in the gut increased as cirrhosis severity worsened. Among salivatypes, we found that SAL1 was enriched with *Prevotella* and *Neisseria* which are known oral commensal bacteria and predominated in HCs. SAL2 conversely was enriched with opportunistic pathobionts, including *Escherichia* and *Campylobacter*, which are typically commensal in the lower intestine and not usually present in the oral cavity. The relative proportion of the more pathogenic SAL2 salivatype - like ENT2 and ENT3 in the gut - also increased as cirrhosis severity and hepatic decompensation worsened (**Figure 1H**). In summary, the fractions of both enterotypes and salivatypes that were enriched with opportunistic pathobionts (ENT2, ENT3 and SAL2) increased significantly with worsening cirrhosis severity.

Additional exploration of the fungal constituents of the microbial communities *-* the mycobiome - demonstrated decreasing diversity (**Figures 1I & 1J**) affecting both the mouth and gut, with compositional changes at phylum level relating to cirrhosis severity (**Supplementary Figure 3**). The gradual increase of Ascomycota with cirrhosis severity led to fungal species level analysis in both oral and gut samples, calculating significance based on Wilcoxon rank-sum test. In the gut mycobiome, *Candida albicans* and *Candida parapsilosis* both significantly increased with cirrhosis severity (**Figures 1K**). In the oral mycobiome, the presence of *Fusarium solani* and other *Fusarium* species significantly increased with cirrhosis severity (**Figure 1L**).

### Overlap between oral and gut microbiome community structures is associated with increasing cirrhosis severity

We demonstrated that pathogenic enterotypes and salivatypes are enriched with bacteria that are usually commensal within a different anatomical niche. For example, *Escherichia* as a commensal intestinal bacteria was enriched in pathogenic salivatype SAL2, whereas commensal oral microbes, such as *Veillonella* and *Streptococcus*, were enriched in pathogenic enterotypes ENT2 and ENT3 (**Figures 1E and 1F**).

These microbes increasingly co-exist and ‘overlap’ in both the oral and gut niches as cirrhosis severity worsens, whilst the degree of overlap in the relative proportions of these different types of bacteria increases, mirroring disease progression (**Figure 3A**). Based on co-existing Metagenomic Species Pan-genomes (MSPs) in both gut and oral sites, including *Streptococcus* spp., *Veillonella* spp., *Escherichia* spp., *Enterococcus* spp., and *Lactobacillus* spp., we identified ordinations of oral and gut metagenome samples (**Figures 1C** and **1D**). Notably, we found that the oral and gut microbiome community structures were more similar to each other as disease severity increased from SC to AD, and to ACLF, in keeping with greater compositional overlap.

We then classified individuals based on the degree of similarity of their oral and gut microbiome into two binary groups: “close” and “distant”, with close describing a higher degree of overlap in bacterial genera between oral and gut niches, and distant being the converse and more in keeping with what is observed in health. Based on this classification, we explored a variety of relevant clinical parameters that might impact upon and/or be affected by the degree of oral and gut microbiome community overlap. These included cirrhosis aetiology, disease severity scores (MELD and Child-Pugh), decompensating symptoms (ascites, HE), ammonia levels, antimicrobial and laxative treatments, gastric acid suppressing treatments (proton pump inhibitor (PPI) and H2 receptor antagonists), non-selective beta-blocker treatments (NSBB) that can affect gut motility, and 1-year mortality (**Figure 3**).

Here we found that worsening disease severity characterised by MELD score and Child Pugh grade (independent of how patients were clinically cohorted in the study) (**Figures 3B and 3C**, respectively), and higher plasma ammonia levels (**Figure 3D**) were observed amongst those cirrhotic patients who had a higher degree of oral-gut microbiome overlap. Alcohol as a cause for cirrhosis was also associated with greater overlap whilst those with metabolic-associated steatotic liver disease (MASLD) were observed to have more distinct oral and gut microbiomes (**Figure 3M**). Notably, drug therapies thought to impact on microbiome composition and some that directly impact upon gut function and the luminal microenvironment, such as antimicrobials, laxatives, gastric acid suppressants and NSBB (**Figures 3E-3I**), were not associated with alterations in the degree of oral and gut microbiome overlap. Specifically, H2-receptor antagonists, whilst trending towards being associated with more of an overlap between oral-gut community structures, did not reach statistical significance (**Figures 3G**).

### Pathogenic entero- and salivatypes are increasingly enriched with virulence factors with worsening cirrhosis severity

We next explored the enterotypes and salivatypes identified based on their putative functional profiles. By aligning the sample-specific gene count profiles with KEGG orthology (KO) annotations, we generated functional profiles summarising all gene counts per KO detected. A total of 10,007 and 15,464 KOs were annotated in all the faecal and saliva samples that were sequenced, respectively.

We first compared KO profiles between pathogenic enterotypes, ENT2 and ENT3, and commensal enterotype, ENT1, and identified 3,072 enriched and 4,429 depleted KOs (Wilcoxon rank sum tests p-values < 0.01). We performed enrichment analysis based on hypergeometric tests and found enriched pathways and modules amongst enriched/depleted KOs in ENT2/ENT3 (hypergeometric tests p-values < 0.01; **Figure 2A** and **Supplemental Figure 1**). Notably, we observed that ENT2/ENT3 harboured genes encoding for the phosphoenolpyruvate:sugar phosphotransferase system (PTS) system which has long been established to have primary functions including sugar transport and phosphorylation as well as sugar reception for chemotactic responses in bacterial cells (*48*). Secondary functions include various ramifications for metabolic and transcriptional regulation with the ability to hijack nutrients from the host.

**Figure 2.**
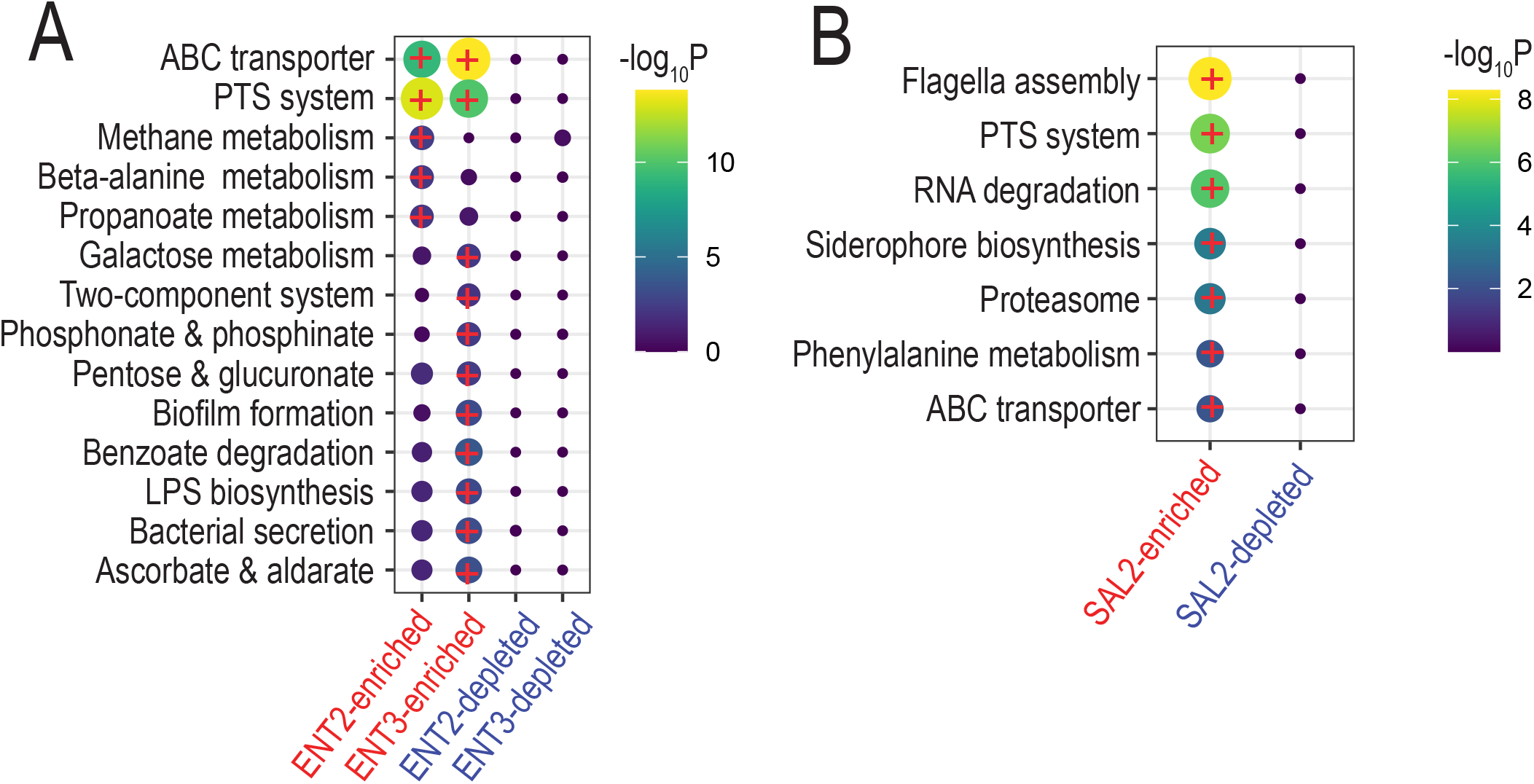
pathogenic enterotypes and salivatypes harboured virulent factors and PTS system hijacking host nutrients. (A)We identified enriched KEGG pathways from enriched/depleted KOs of pathogenic enterotypes, ENT2 and ENT3, compared to ENT1 (hypergeometric tests p-values < 0.01). (B) We also identified enriched KEGG pathways from enriched/depleted KOs of pathogenic salivatype, SAL1, compared to SAL1 (hypergeometric tests p-values < 0.01).

**Figure 3.**
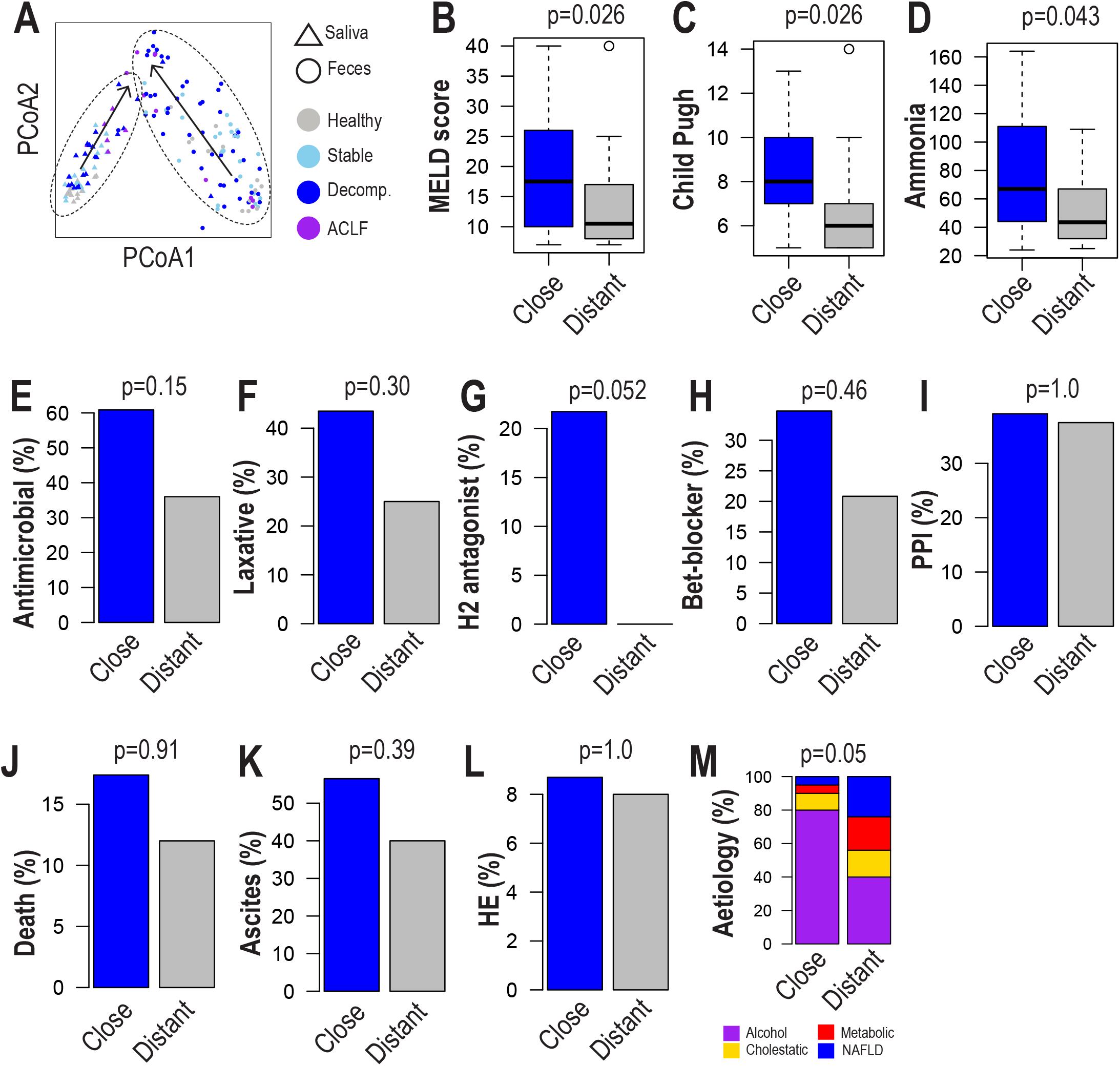
gut-oral translocation linked to disease severities. A, ordination plots of Bray-Curtis dissimilarity of faeces and saliva samples based on translocating species between gut and oral sites. We found that gut and oral microbiome tended to be similar while severity increases (see arrows). B-O, we grouped subjects based on similarity of gut and oral microbiome – “close” and “distant” groups. We identified that disease severity and other pathogenic parameters increase among “close” subjects. B: MELD score, C: Child Pugh score, D: Ammonia level, E: antimicrobial treatment (%), F: laxative treatment (%), G: H2 receptor antagonist treatment (%), H: beta-blocker treatment (%), I: proton pump inhibitor (PPI) treatment (%), J: death at 1 year (%), K: Ascites (%), L: hepatic encephalopathy (HE) (%), M: aetiology. Wilcoxon rank tests and chi-square tests were performed for continuous and categorical variables, respectively.

We also found that bacterial genera within ENT3 harboured more virulence factors, including biofilm formation (*49*), lipopolysaccharide (LPS) biosynthesis (*50*), bacterial secretion systems (*51*), and ascorbate degradation, which can initiate inflammation, transfer virulence factors and hijack host nutrients. Dissimilatory nitrate reduction modules that generate ammonia - central to the pathogenesis of HE - were additionally found to be enriched in advanced cirrhosis patients harbouring ENT2 and ENT3 in the gut. Biofilms promote horizontal gene transfer through the exchange of bacterial genome fragments and/or mobile genetic elements, which contributes to the spread of antibiotic-resistance genes (*52*).

We also compared KO profiles between SAL2, pathogenic salivatype, and SAL1, commensal salivatype (Wilcoxon rank sum tests p-values < 0.01). Among 2,503 enriched and 5,980 depleted KOs in SAL2 samples, we identified enriched pathways and modules based on hypergeometric tests (p-values < 0.01; **Figure 2B and Supple Figure 2**). Here we also identified many virulence factors that can contribute to pathogenic properties of those bacteria in SAL2, including flagella assembly, siderophore biosynthesis, PTS system, autoinducer (AI)-2 transport, and type IV secretion system (*53*).

The phosphotransferase system (PTS) is one of the sugar transport systems in oral microbes. PTS can transport galactose and produce galactose 6-phosphate, whilst ornithine, ammonia, and carbon dioxide are generated by enzymatic degradation of citrulline and ammonia by arginine deiminase. This series of reactions referred to as the arginine deiminase system are considered to have developed to oppose sugar metabolism-based acidification (*54*). Here we found the up-regulation of carbohydrate transport and metabolism in both oral (SAL2) and gut (ENT2, ENT3) microbiome of the sicker cirrhotic patients, including significant enrichment of PTS system, a sugar transport system that hijack host nutrient and also enrichment of galactose metabolism, of which excess metabolites, such as galacitol can lead to oxidative stress or act as metabotoxin (*55*).

In summary, not only are there significant alterations in microbial community structures in the oral and gut compartments in cirrhosis which become more pronounced as disease severity progresses, there are several putative functional alterations in these altered microbial communities which have multiple pathogenic properties that mirror disease progression and which have the potential to impact deleteriously on the host.

### Alterations in oral and gut microbiome antimicrobial resistance genetic profiles

To investigate the frequency and potential for harbouring of antimicrobial resistance genes (ARGs) in this study, we profiled ARGs within the oral and gut microbial datasets utilising the Comprehensive Antimicrobial Resistance gene Database (CARD) database (**Figure 4A**). We found that in the majority of individuals, the oral and gut microbiome harboured substantial numbers of ARGs (1,218 and 672 genes for oral and gut microbiome, respectively). Many of these ARGs were common to both sites (575 genes), although a greater proportion of these shared ARGs were detected in the gut (>85%) compared to the oral niche (47%). We then checked whether total ARG abundances of the oral and gut microbiome differed based on CLD severity and compared to the HC and NLS cohorts (**Figure 4B and C**). Notably, total ARG abundances in both the oral and gut microbiome increased as CLD severity and hepatic decompensation worsened, with this pattern more pronounced for the gut resistome, as has been previously hypothesised. For both niches, the NLS group demonstrated highest total ARG abundances.

**Figure 4.**
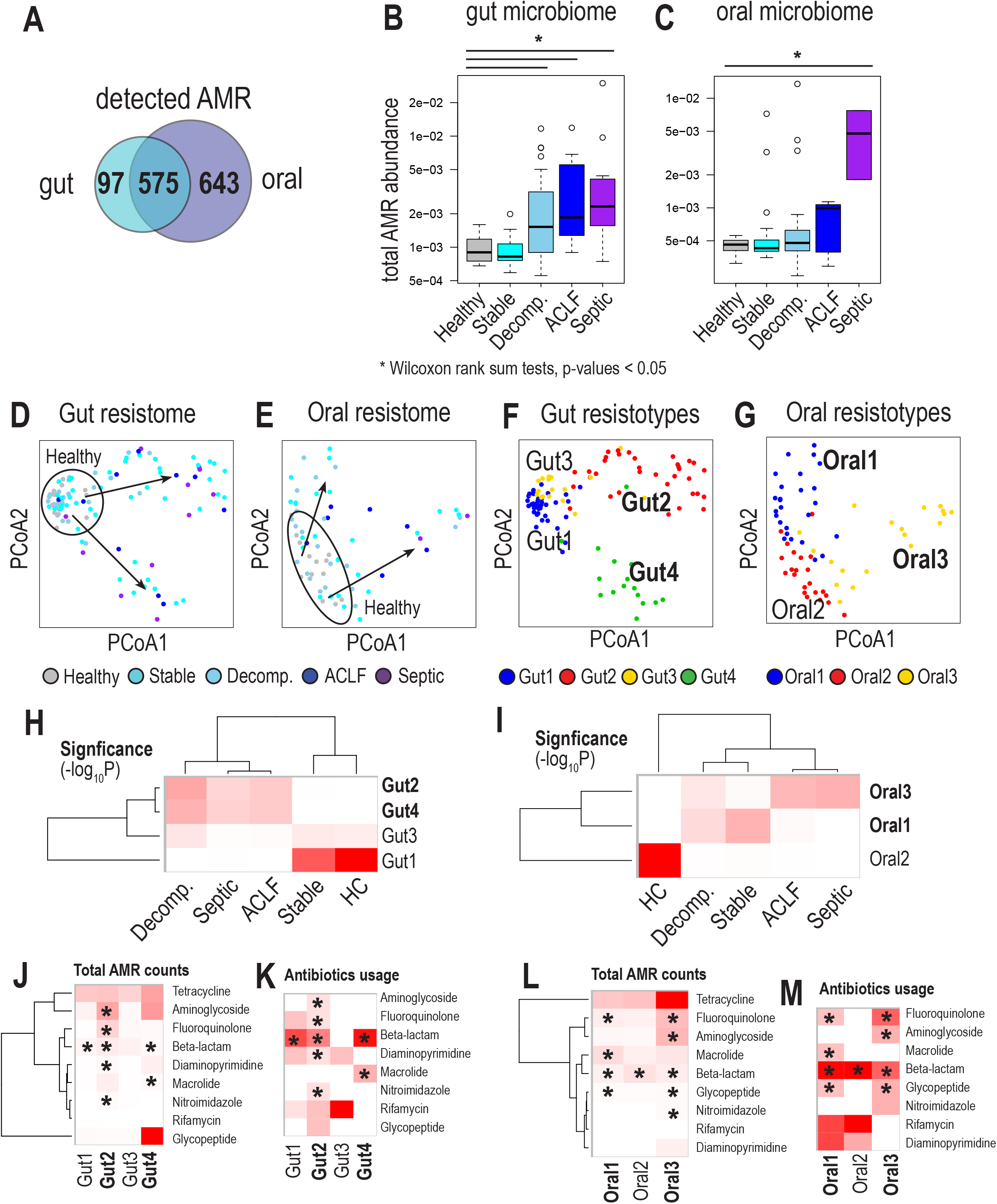
Distribution of antimicrobial resistant genes among cirrhosis patients. (A) we identified antimicrobial resistant (AMR) genes in faecal and saliva shotgun metagenomic samples and observed the majority of them were shared between faecal and saliva samples. We estimated total gene abundance of all AMR detected in (B) faecal and (C) saliva samples and compared them by disease severity. Enhanced AMR abundance was observed with increasing disease severity (Wilcoxon rank sum tests p-values < 0.05). To further explore AMR classes enriched with different severities, we performed principal coordinate analysis of AMR profiles of (D) gut and (E) oral microbiomes. We found that healthy samples were clustered separately from samples from cirrhotic patients.. We identified resistotypes by performing Dirichlet multinomial mixture modelling from (F) faecal and (G) saliva samples – Gut1/2/3/4 and Oral1/2/3. (H) Gut1 and Gut3 were enriched among healthy and stable cirrhotic subjects, whereas Gut2 and Gut3 were enriched amongst the sicker cirrhotic patients. (I) Oral1 and Oral3 were enriched in those with decompensated cirrhosis, whereas Oral2 was enriched in healthy subjects. We checked (J) AMR abundances of gut resistotypes and (K) fractions of subjects with antibiotic prescriptions, per antibiotics class. We also checked (L) AMR abundances of oral resistotypes and (M) fractions of subjects with antibiotic prescriptions, per antibiotics class. We found that AMR genes for beta-lactam antibiotics were spread in the majority of gut and oral resistotypes. Star marks indicate simultaneous observed antibiotics classes with high AMR abundance and antibiotics subscriptions.

To further explore the ARG profiles (resistome) of oral and gut samples across the different CLD severities, we performed principal coordinate analysis (PCA) of ARG profiles for both niches (**Figure 4D and E**). As previously described, we found that HCs harboured unique oral and gut microbiome resistomes. However, the resistome of cirrhotic patients was significantly different to that of the HCs. By performing unsupervised clustering, three and four resistotypes (*56, 57*) or the oral and gut microbiome, respectively, were determined (**Figure 4F and G**). Of the gut resistotypes, Gut1 and to a lesser extent Gut3 were both enriched amongst the HC and SC cohorts, in contrast to Gut2 and Gut4 which were enriched in patients with AD and ACLF and to a lesser extent in NLS (**Figure 4H**). For the oral resistotypes, Oral2 was specific to HCs whilst Oral1 was most enriched in SC and then AD, whilst Oral3 was most enriched in ACLF and NLS and then in AD.

ARG classes were assessed for oral and gut resistomes, based on their drug classifications as determined by CARD. We detected enrichment of ARGs for β-lactamase classes in all oral and all but one of the gut resistotypes. ARGs encoding for resistance against aminoglycoside, fluoroquinolone, macrolide and nitroimidazole drug classes were specifically enriched in oral and gut resistotypes specific to AD and ACLF patients, including Oral1/Oral3 and Gut2/Gut4 (**Figures 4J and 4L**). We observed that there was a high proportion of antibiotic use in cirrhosis patients, where their oral and gut resistotypes indicated resistance against those specific types of antibiotics that patients were treated with (starred in **Figures 4K and 4M**). A high degree of ARGs to β-lactamase inhibitors (e.g. piperacillin-tazobactam) and carbapenems (*e.g.* meropenem) were detected in all but the Gut3 resistotype in AD and ACLF patients, with 87.9% of all these patients receiving some form of β-lactamase antibiotic; 71.4% and 16.5% were simultaneously being treated with either a β-lactamase inhibitor or a carbapenem, respectively.

ARGs for rifamycin from which rifaximin-ɑ, a prophylactic therapy used in HE, is derived, were not significantly increased in abundance in the majority of cirrhosis patient oral and gut samples. This is despite up to 25% of SC, AD and ACLF patients being either concomitantly or treated up to hospital admission with rifaximin-ɑ for secondary prophylaxis for HE.

## Discussion

In this study we performed in-depth shotgun metagenomics to elucidate the alterations of oral and gut microbiome compositions and, crucially, interrogated aspects of specific functional alterations in distinct severities of cirrhosis. This was achieved by the simultaneous assessment of bacterial and fungal components of the saliva and faecal microbiome, as surrogates for the oral and gut environments, in a robustly phenotyped cohorts of cirrhosis patients. These findings have been contrasted with healthy individuals and uniquely to a positive control cohort of patients with sepsis but without underlying cirrhosis. After interrogating the community structures, we provide additional novelty by evaluating virulence factors that provide insight into the putative functions of the oral and gut microbiome, as well as assess ARG abundance based on oral and gut ‘resistotypes’, and how the so-called resistome alters as cirrhosis severity progresses and relates to antimicrobial exposure.

We identified simultaneous substantial bacterial and fungal alterations in the gut and oral microbiome, beginning with a significant reduction in (alpha-)diversity affecting both communities as cirrhosis progresses. This is consistent with previous reports where one or other gut/oral microbial community has been studied, in relation to decompensation with HE, pharmacotherapies such as PPI, and/or hospitalisation (*16, 58*). Recent large cohort studies have reported on the utility of simultaneous evaluation of salivary and faecal microbiome in cirrhosis (*59*) and when comparing cirrhotic cohorts from the USA and Mexico, where greater linkages between the faecal microbiome with plasma metabolites, compared to saliva, were reported (*60*). These studies were however limited by employing lower resolution V1-V2 16S rRNA gene analysis instead of V3-V4 analysis or deep metagenomics used in this study.

Family-level alterations affecting both the oral and gut microbiome as cirrhosis severity worsened showed that opportunistic pathobionts (*Enterococcaceae*, *Enterobacteriaceae*, *Veillonellaceae* and *Streptococcaceae*) were over-represented in both anatomical niches. In contrast, the reduction in relative abundance of indigenous bacteria in the mouth (*Neisseriaceae* and *Prevotellaceae*) and the gut (*Oscillospiraceae* and *Ruminococcaceae*) with worsening cirrhosis has implications for host-requiring metabolic activities including nitrate reduction and butyrate production, respectively. Whilst previous studies have focused more on gut alterations, our findings are also consistent with oral microbiome studies whereby *Veillonella* was associated with cirrhosis and *Nesseria* associated with healthy individuals (*61*). Functional prediction in this study demonstrated a significantly higher proportion of genes associated with carbohydrate transport and metabolism, defence mechanisms and membrane transport, all indicative of enhanced pathogenicity, and mirroring our data. Our findings have implications for interactions of these communities along the oro-intestinal tract and altered functional relationships affecting the human host in cirrhosis.

In addition to the evolving concepts of ‘invasion’ and ‘oralisation’ of the intestinal microbiome in CLD(*62*), there is now considerable focus on the role of the oral microbiome which is increasingly recognised as predisposing to hepatic decompensation (*16, 63*), in addition to the gut microbiome. In this study, commensal enterotype (ENT1) and salivatype (SAL1), which were enriched e.g. *Oscillospiraceae*, and *Prevotellaceae*, were found to be significantly reduced, whereas pathogenic enterotypes (ENT2 and ENT3) and salivatype (SAL2) were significantly increased. Substantial overlap of gut and oral microbiome communities, such as *Enterococcaceae*, *Streptococcaceae*, and *Veillonellaceae*, for both pathogenic enterotype and salivatype, may imply bi-directional colonisation from not only the oral to more distal intestinal niches, but also from intestine to the more proximal oral niche. The relocation of bacteria from the oral to intestinal niche has been reported in advanced cirrhosis (*27*). Rifaximin-α was reported to suppress the growth of orally originating species - commonly found in dental plaque and associated with periodontal disease - in cirrhotic faeces in the setting of a randomised controlled trial. These oral species have putative functions related to intestinal mucus degradation, such that a reduction in these pathobionts seeding into the gut promotes gut barrier repair, further emphasising the intimate relationship of the oral-gut-liver axis.

Alterations in functional capacity of pathogenic enterotypes and salivatypes implicate changes in the homeostatic metabolism of gut and oral microbiota, providing new opportunities for pathobionts to disrupt homeostatic mechanisms by several processes. Firstly, host nutrient hijacking by PTS system, ABC transporters, siderophore biosynthesis, and ascorbate degradation (*64, 65*); (2) host tissue invasion due to enhanced bacterial secretion systems and flagella assembly (*66*); (3) promoting host inflammation by LPS biosynthesis (*67*) and (4) promoting dysfunctional metabolic pathways that generate greater oxidative stress by pentose and glucuronate metabolism (*68*), and galactose metabolism (*69*). In relation to ammonia metabolism, dissimilatory nitrate reduction modules that generate ammonia were enriched in advanced cirrhosis, implicating increased ammonia production in AD and ACLF patients. Ammonia is central to the pathogenesis of hepatic encephalopathy (HE) (*70*) and may causally link these changes in gut microbiome putative metabolic functions with driving the development of HE in cirrhosis (*11*).

The profile of ARGs within a microbial community is known as the ‘resistome’ (*71–73*). Exploration of the resistome by NGS provides valuable insight into mechanisms behind development of MDROs (*74*). ARGs can also represent quorum-sensing and secretion system survival strategies independent to antibiotic exposure (*75*), regulate ecological dynamics within an environment, and determine survival in complex microbial communities due to adaptations in phenotypic and genotypic responses to antimicrobials (*76*). In common with our previous work (*77*). We identified that there are key differences in the overall resistome profile between the gut and oral cavity, with different resistor types in each site. In addition, the lack of a significant rise in total ARG abundance in the oral microbiome in contrast to the gut microbiome supports the idea that the oral resistome is inherently more stable, as has been proposed by us and others (*77, 78*). As part of the functional analysis, striking alterations in the ARG repertoire were identified in both gut and oral microbiome, based on cirrhosis severity. However, it was notable that resistance to the most commonly used antibiotic class, the β-lactams, did not show a similar level of increase. Whilst this is potentially due to the increased resistance of piperacillin to Gram-negative β-lactamases, it is more likely due to the use of combination drugs, such as piperacillin/tazobactam. In these instances, the co-use of tazobactam (or similar compounds) will inhibit the action of the bacterial β-lactamases (*79*), extending the efficacy of the antibiotic and suppressing the development of resistance by counteracting the beneficial (to the microbe) impact of β-lactamases.

The alterations described here of the oral and gut resistome appear to reflect disease type and severity. Another study comparing faecal ARG burden between compensated and DC patients with hepatic encephalopathy (HE) and ascites, reported high gut ARG counts in cirrhosis (*59*) and interestingly was distinct from the resistome in chronic kidney disease and diabetes mellitus. Faecal ARG burden worsened with disease progression regardless of ascites and HE and was associated with risk of hospitalisation and death, independent of cirrhosis severity, prior antibiotic exposure, hospitalisations, or concomitant medications. We have previously demonstrated that there are also key differences between the oral resistome and the gut resistome in healthy individuals, demonstrating reduced diversity but increased abundance of ARGs in the oral cavity, relative to the gut (*77*). This study is the first to describe and contrast in detail the oral and gut cirrhotic resistome simultaneously, and how this changes as disease severity progresses. These data also indicate that whilst ARGs may be more positively selected based on antibiotic exposure, there are potentially additional selection pressures to select for these ARGs beyond direct antibiotic-induced selection pressure, with the most significant increases in ARGs not associated with treatment, such as tetracycline.

We have previously shown that biomarkers of intestinal inflammation and barrier damage rise with increasing cirrhosis severity (*80*). Increased ARG carriage in patients with cirrhosis may be driven by intestinal inflammation and gut barrier damage (*81*). Previous studies have linked gut inflammation with enhanced AMR gene transfer, potentially through a rise in horizontal gene transfer (*82*). It is possible that these ARGs are co-selected with other genes, such as those involved in virulence, or that they are carried at higher levels in pathobionts identified in the context of loss of bacterial diversity as cirrhosis progresses. It is important to note, however, that we are unable to ascertain what is cause and effect in those sicker patients more frequently treated with antibiotics. Based on the discovered resistotypes and further studies, it may be possible to begin to tailor antibiotics in cirrhosis to avoid selecting for AMR dominant microbes, and thereby improve pharmacotherapeutic effectiveness.

By utilising whole-genome shotgun sequencing, we were able to simultaneously assess the gut and oral mycobiome in addition to the bacterial communities, and report novel findings in this cirrhotic population. The understanding of how microbiome alterations relate to disease in general have traditionally been largely bacteria- and gut-focused (*83*), but emerging evidence suggests that fungi also play an important role (*84*). Fungal communities represent a relatively minor component (0.03%-2% of the total) of the gut microbiome, compositionally (*85*) and mycobiome exploration remains in its infancy, and even more so in the context of cirrhosis. Offering additional insights into non-bacterial communities, we detected an overall reduction in fungal diversity affecting both oral and gut niches as cirrhosis severity worsened, and which was more pronounced in the intestine, mirroring the bacterial community alterations. In tandem, however, certain fungal genera such as Candida and Fusarium increased in relative abundance in the gut and mouth, respectively, with increasing cirrhosis severity.

There is increasing recognition that the gut mycobiome can impact on cirrhosis pathogenesis and severity and associate with different aetiologies (*86, 87*). The literature is more sparse around the role of the oral mycobiome in liver disease *per se*. In cirrhotic faeces at least, mycobiome alterations affecting *Candida* have been reported that alter with antibiotic and PPI use (*88*). The gut mycobiome is increasingly considered as a central component of the microbiome overall in cirrhosis (*86*), either as a consequence of the disease process or having a pathogenic contribution in aetiology and progression. *Candida albicans* has been reported to be of higher abundance in the gut of patients with ArLD and MASLD (*89*) whilst *Candida parapsilosis* was higher in the faeces of hepatitis B cirrhosis patients compared to healthy volunteers (*90*). From a functional perspective, *C. albicans*-produced exotoxin candidalysin promotes ArLD in preclinical models (*91*) whilst *C. parapsilosis* shows an increasing abundance of antifungal resistance genes (*92*), both emphasising the pathogenic role of the mycobiome in CLD.

The finding of Fusarium in the oral mycobiome to cirrhosis is a novel finding, with an increase in abundance as disease severity worsens. Mould-like species such as *F. solani* have been reported in ArLD but are confined to the gut (*93*). Fusarium has also been associated with a variety of invasive diseases and as a mould, the hyphal filaments are efficient in breaching an already dysfunctional intestinal barrier (*94*). Immune deficiency, antibiotic use and indwelling medical devices may increase the risk of invasion by mould-like fungi such as Fusarium *(95)*. Fusarium is known to be invasive in immunodeficient disease states such as haematological cancers (*96*). With limited information about the Fusarium genus in cirrhosis, there is accumulating evidence for the role of intestinal mycobiota in liver disease. ArLD with hepatocellular inflammation displays overgrowth of invasive intestinal fungi such as C. albicans, further supported by the presence of ß-glucan fungal cell wall component in plasma (*97, 98*). Faecal mycobiome composition varies based on severity of MASLD, with elevation of plasma anti-C. albicans IgG levels in advanced fibrosis. Previous liver disease studies have focused mainly on bacteriome and fungal profiles at phylum and genus phylogenetic levels. Mycobiome studies have only confirmed similar findings of reduced fungal diversity that is thought to be related to the use of antibiotics causing reduction of microbiota diversity (*99*). It is possible that antibiotic-induced dysbiosis encourages the outgrowth of invasive fungal species. The direct cause-and-effect relationship between the mycobiome and cirrhosis remains to be determined but evidence suggests that fungal products contribute to hepatocyte damage with the promotion of an inflammatory response (*97*). Nevertheless, observations made here and previous studies confirms the significance of gut mycobiome in liver disease and need for further evaluation (*100*).

In view of the single centre, largely Westernised cohort of cirrhosis patients and controls employing single time point biological sampling, future work to expand on these findings will require a multi-centre approach that involves patients from a wider spectrum of ethnicities and backgrounds. Whilst there are advantages to a cross-sectional approach to determine differences in microbial communities between different anatomical niches and patient cohorts, longitudinal studies are now required to determine how microbiome changes alter as cirrhosis progresses and complications occur, to begin to better evaluate causality and provide a more comprehensive perspective on microbiota diversity (within-subject and between-subject diversities) (*101*) as well as develop more sophisticated therapies. In addition, integrating these high resolution metagenomic datasets with more detailed oral health assessment, nutritional and other environmental and lifestyle factors will allow better understanding of the effect of these potential confounders (*102*) in addition to the impact of disease severity and phenotype (18).

Therapeutically targeting the gut microbiome is increasingly drawing attention as a potential approach in cirrhosis (*103–105*) as well as for specific complications such as(*106, 107*) HE, as knowledge expands around the putative role of the alterations to which this study contributes. Therapies being proposed vary from relatively untargeted approaches utilising pre-biotics and probiotics, nanoporous carbons, farnesoid X receptor agonists and faecal microbiota transplantation, to more precision-based therapies, including engineered bacterial strains, phages and postbiotics (14). This is especially relevant in an era of increasing AMR infections in cirrhosis, where non-antibiotic dependent approaches need to be developed (7). In addition to the gut, the oral microbiome is now also considered in cirrhosis to be targetable by periodontal therapies as well as nutrition-based approaches in cirrhosis (*108, 109*). In order to realise the full potential of such approaches whilst minimising therapeutic misadventures, and to determine which strategies are best for individual patients, enhancing our fundamental understanding of oral and gut microbiome functional alterations remains crucial in cirrhosis to fully exploit these pathways.

**Supplementary Table 1.** Presence of bacterial infections among different clinical groups.

| **Groups** | **SC** | **DC** | **ACLF** | **SEP** |
| --- | --- | --- | --- | --- |
| **Blood culture** | 3 | 16 | 3 | 5 |
| **Sputum culture** | 2 | 4 | 2 | 3 |
| **Urine culture** | 4 | 9 | 2 | 3 |
| **Ascites WBC** | 3 | 10 | 7 | 1 |
| **Beta D-glucan** | 0 | 1 | 4 | 0 |
| **MDRO** | 0 | 5 | 0 | 1 |

**Supplementary Figure 1.** Enriched biochemical pathways of enterotypes. We identified enriched KEGG modules from enriched/depleted KOs of pathogenic enterotypes, ENT2 and ENT3, compared to ENT1 (hypergeometric tests p-values < 0.01).

**Supplementary Figure 2.** Enriched biochemical pathways of salivatypes. We identified enriched KEGG modules from enriched/depleted KOs of pathogenic salivatype, SAL1, compared to SAL1 (hypergeometric tests p-values < 0.01).

**Supplementary Figure 3.** Relative abundance of the fungal phylum.

## Acknowledgements

We are grateful to all the patient participants and healthy volunteers for agreeing to take part in this study, and to the clinical and liver research teams at King’s College Hospital for facilitating recruitment, collecting metadata and sample collection. We thank the King’s College Hospital Institute of Liver Studies and Transplantation Charitable Research Fund and a generous donation from a Liver Service User, and the Foundation for Liver Research for funding and facilitating this work within The Roger Williams Institute of Hepatology.

We acknowledge support from the National Genomics Infrastructure in Stockholm funded by Science for Life Laboratory, and SNIC/Uppsala Multidisciplinary Center for Advanced Computational Science for assistance with massively parallel sequencing and access to the UPPMAX computational infrastructure, under project number Project SNIC 2020-5-222, SNIC 2019/3-226, SNIC 2020/6-153, SNIC 2021/6-89, SNIC 2021/5-248 and SNIC 2021/6-242.

## Data Access

The shotgun metagenome raw data, sequenced as part of this study can be found from EBI ENA repository under the project accession PRJEB52891.

