## Supplementary figures and images for "Pathogenic entero- and salivatypes harbour changes in microbiome virulence and antimicrobial resistance genes with increasing chronic liver disease severity"

### Supplementary Figure 1

##### Supple Figure 1

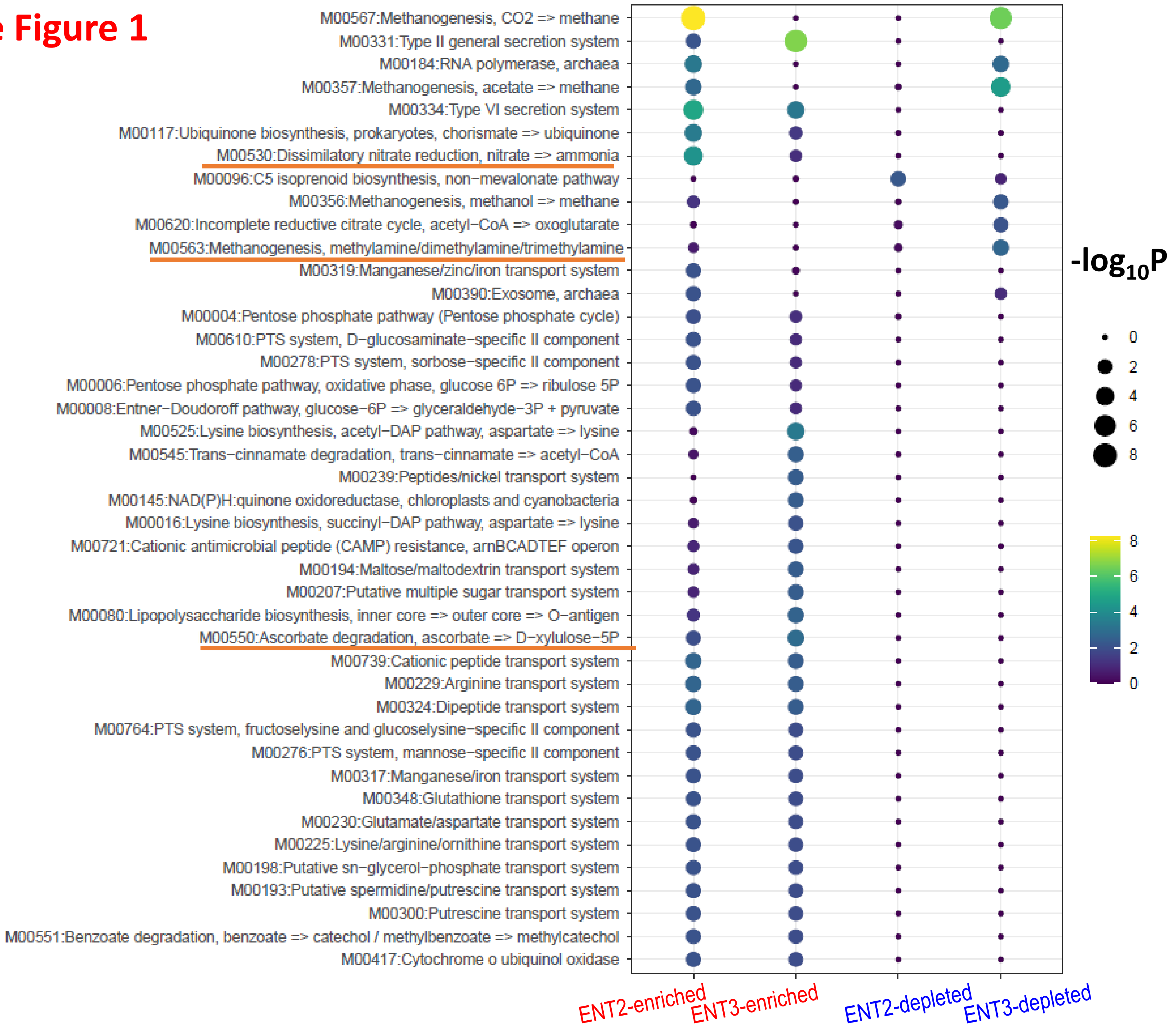

### Supplementary Figure 2

#### Supple Figure 2

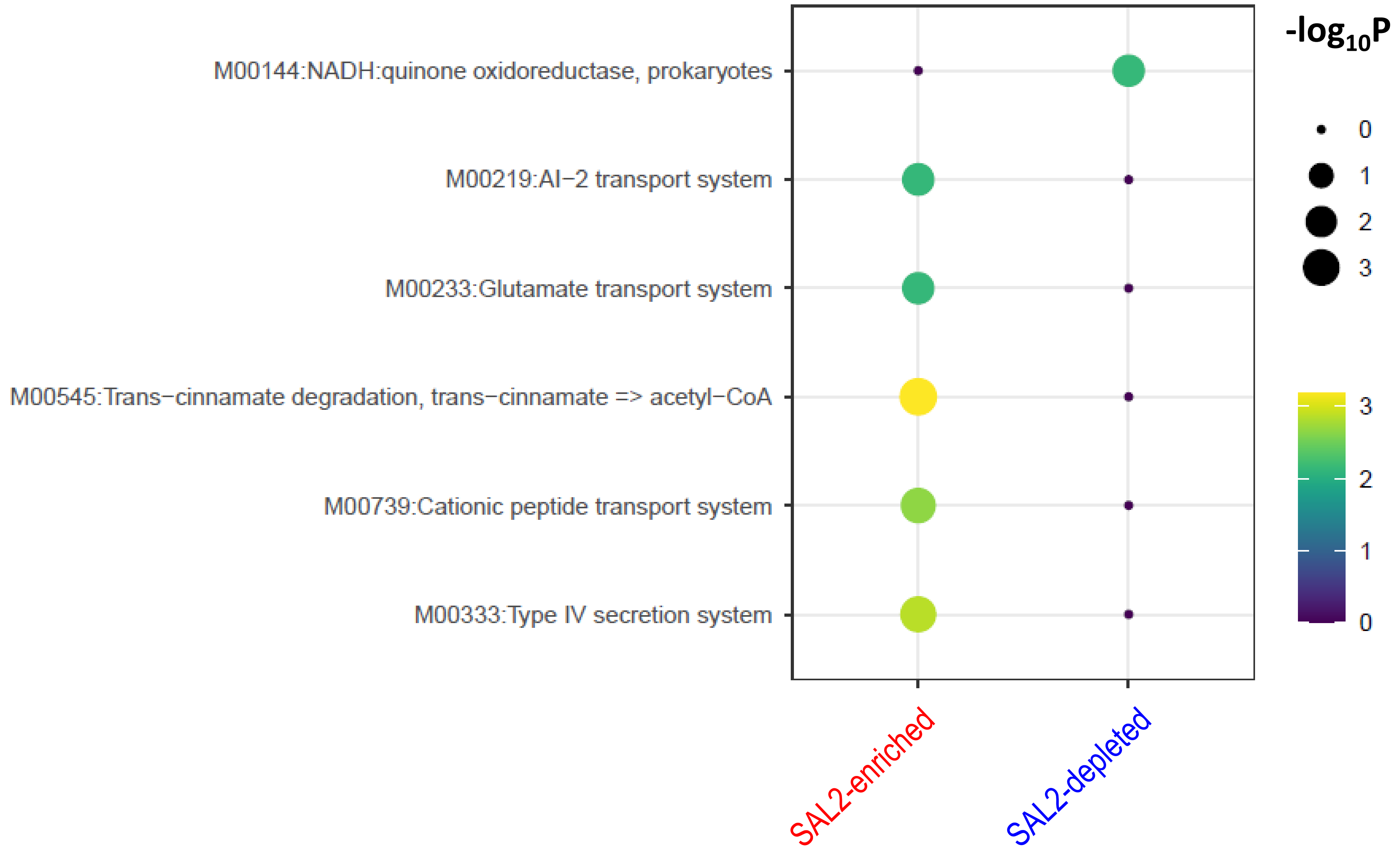

### Supplementary Figure 3

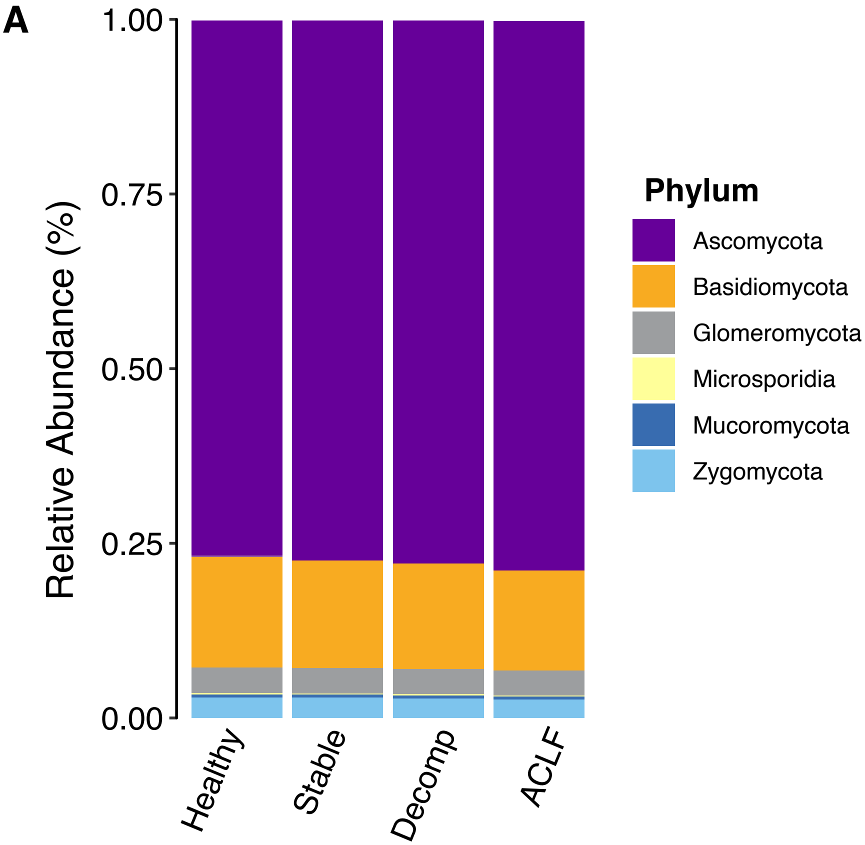
